# Novel Smartphone Based Free Flap Monitoring Tool Using Machine Learning

**DOI:** 10.1101/2022.07.28.501864

**Authors:** Destie Provenzano, Akash Chandawarkar, Edward Caterson

## Abstract

Free flap monitoring is important to ensure early detection of arterial or venous failure to facilitate salvage. Our prior research has shown ability to magnify skin change as a result of skin changes. This study was undertaken to test the feasibility of detecting venous and arterial occlusion using a smartphone camera and pattern recognition (a simplistic implementation of a machine learning algorithm). Bilateral hands of seven patients were video recorded with various tourniquet pressures on one hand simulating no occlusion, venous occlusion, and arterial occlusion with the other hand as internal control. Video data resolved at an average iPhone camera quality of 33 fps was processed using the sci-kit learn library in Python to detect changes in color frequency between frames and then compared to the control hand. Comparing the test hand to the control hand allowed for the depiction of the “delta” that was sensitive enough to detect changes on a video without any additional augmentation. The average rate of change in red pixels between video frames was noticeably different compared to control for both arterial occlusion (1.06x greater) and venous occlusion (1.07x greater). A graphical representation depicted a clear relationship while an individual was undergoing occlusion (Fig 1). Our smartphone video capture and analysis facilitates visualization of skin perfusion and can distinguish between states of no occlusion, arterial occlusion, and venous occlusion. This study shows promise for the use of inexpensive smartphone monitoring in a clinical setting for accurate free flap monitoring.

## I. Introduction

Maintenance of vascular sufficiency is fundamental to the ultimate success of any free flap transfer. Among other complications, postoperative total flap failure can require salvage surgery, increase hospitalization time, and ultimately decrease patient quality of life. [1,2] It’s important to detect vascular insufficiency in the flap early to facilitate salvage and deter further complications. [3,4] Studies have shown that early detection of vascular insufficiency greatly increases the likelihood of salvage should complications occur. [5-8]

**Figure 1:**
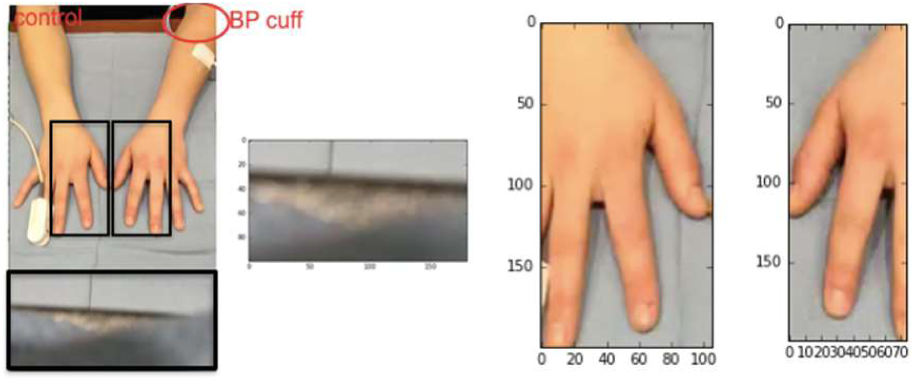
Sample depictions of frame from the video and corresponding cropped regions of a control region, the contralateral hand (leftmost image of hand), and hand undergoing occlusion (rightmost image of hand).

Both microsurgery and the monitoring techniques to evaluate postoperative performance have progressed steadily over the past 30 years with many advances in both skills and technologies. [9-11] Many of these technologies strive to meet the criteria as defined by Creech and Miller. Ideal Free Flap Monitoring systems were defined by Creech and Miller to be “Continuous, Non-Invasive, Direct, Easy to Interpret, Quantitative, and Not Expensive.” [12] Currently no system available on the market meets all of these criteria, as many that are non-invasive and easy to use are also incredibly costly. [13,14]

Our prior research has shown ability to magnify skin change as a result of skin changes. Eulerian magnification, near-infrared spectroscopy, and white light spectroscopy have emerged as promising technologies for post-operative monitoring. [15-17] We previously used Eulerian magnification on simulated instances of arterial and venous occlusion to determine the feasibility of using a camera based monitoring device to detect insufficient perfusion. [18] Although the technology worked, it proved computationally intensive. This study was undertaken to test the feasibility of detecting venous and arterial occlusion using a smartphone camera and pattern recognition (a simplistic implementation of a machine learning algorithm).

We utilized data from seven patients on various scales of the Fitzpatrick skin type spectrum and simulated venous and arterial occlusion using a blood pressure tourniquet. A simplistic pattern recognition algorithm was developed on said data and successfully was able to predict when instances of venous and arterial occlusion was beginning to occur. This pilot study is the first of its kind to showcase a free flap monitoring technology that is cost effective, non-invasive, direct, easy to interpret, quantitative, continuous and not computationally intensive.

## II. Methods

### A. Data Collection

Bilateral hands of seven patients were video recorded with various tourniquet pressures on one hand simulating no occlusion, venous occlusion, and arterial occlusion with the other hand as internal control. The contralateral hand functioned as an internal control.

### B. Data Processing

Video data resolved at an average iPhone camera quality of 33 fps was processed using the sci-kit learn library in Python to detect changes in color frequency between frames and then compared to the control hand. To standardize the “hand” across each video, the initial frame was cropped to each hand and a separate control region of the video (Figure 1).

Comparing the test hand to the control hand allowed for the depiction of the “delta” that was sensitive enough to detect changes on a video without any additional augmentation.

### C. Data Analysis

Video data was processed in the sci-kit learn python library at 33 fps to mirror data captured from a phone camera. The data was processed in a four step process (Figure 2) that worked by breaking each frame into a matrix of color pixels (1), comparing each frame to the subsequent frame in the video to find the “delta” between frames (2), overlaying this delta over a control region of the video to standardize against lighting (3), and then overlaying the affected hand over the control region to control against skin tone and individual fluctuations (4).

**Figure 2:**
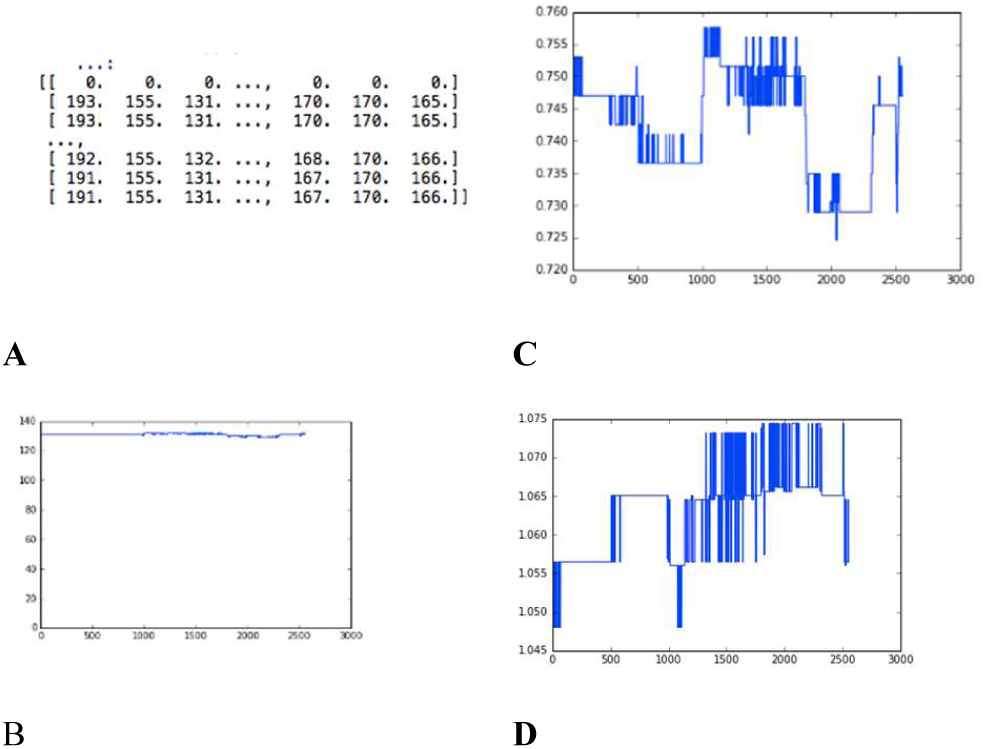
Depiction of data process to calculate the change in perfusion between frames and between hands. A) Each frame of video was broken into a matrix of average color/ row and averaged across each frame. B) Subsequent frames were compared to look for “changes” in increase or decrease of pixel count. C) Occluded hand was overlayed over a control segment of the video to account for changes in lighting and then compared to the control hand. D) Affected hand was overlayed over the control hand.

The average color, total color, and average row color for each frame of the video data was processed and compared frame by frame at 33 bps (33 frames/ second). Pulse was obtained by normalizing the data for changes in the average red color between frames to be > 1.005 for any 2 or more frames in a row. To account for potential changes in lighting and quality of video data, occlusion was determined by overlaying pulse from the occluded hand over the non-occluded hand.

### D. Data Validation (Pulse)

Pulse was calculated using this method for a set of control videos to validate this process. Separate test videos were taken on non-occluded skin in outdoor, indoor, bright, dim, partially shaded, and normal lighting. Additional test videos were taken while walking for approximately one mile outdoors to test if motion affected pulse collection data as well. Several hundred videos of each were recorded across three subjects for one minute at a time on an Iphone 8 camera at 33 fps. Pulse was recorded manually for each attempt by standard methods (A finger was placed on the wrist and beats were counted for one minute while filming).

Each video was processed in the scikit learn python library. Data was first cropped to the region of skin on the hand and overlaid over a control region of the video. The average number of red pixels per frame was calculated and then subtracted between frames to uncover the pulse per minute. The concordance of measure heart rate (pulse ox) and calculated heart rate using the algorithm were used to validate the results of this study.

## III. Results

### A. Pulse Validation Results

The pattern recognition technique seeking to find the delta between frames worked across the control videos and was able to consistently detect pulse within 5 bpm (at a p < 0.01). (Figure 3)

**Figure 3:**
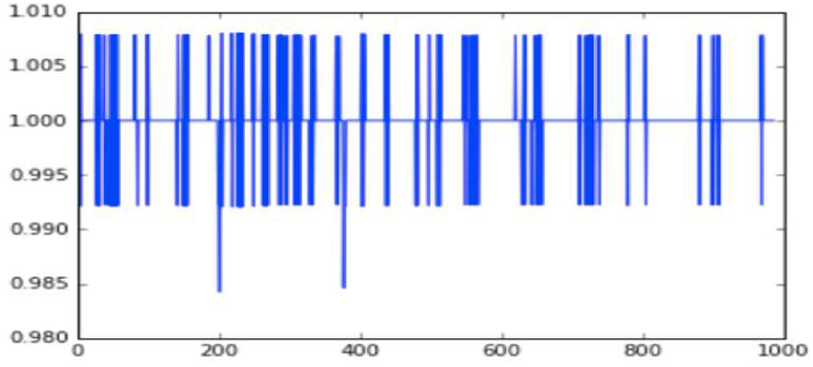
Output from 30 seconds of recording showcasing a heart rate of “70 beats/ min” for non-occluded hand. This depicts the change in pixels between frames. Several hundred control videos of non-occluded skin were used to create the final algorithm. This proved a concordance of measured HR (pulse ox) and calculated HR with algorithm.

It was found that any delta between frames that showed a consistent increase in red pixels (Of non occluded skin with a control region subtracted out to account for lighting) for 2 or more frames corresponded to a heart beat. This method produced accurate measurements of pulse within 5 bpm across all variations in the test variables including movement, lighting, or skin tone.

### B. Occlusion Measurement Results

The average rate of change in red pixels between video frames was noticeably different compared to control for both arterial occlusion (1.06x greater) and venous occlusion (1.07x greater). A graphical representation depicted a clear relationship while an individual was undergoing occlusion (Figure 4). The processed change in average red pixels per frame for the occluded hand overlaid over the control hand are depicted below.

**Figure 4:**
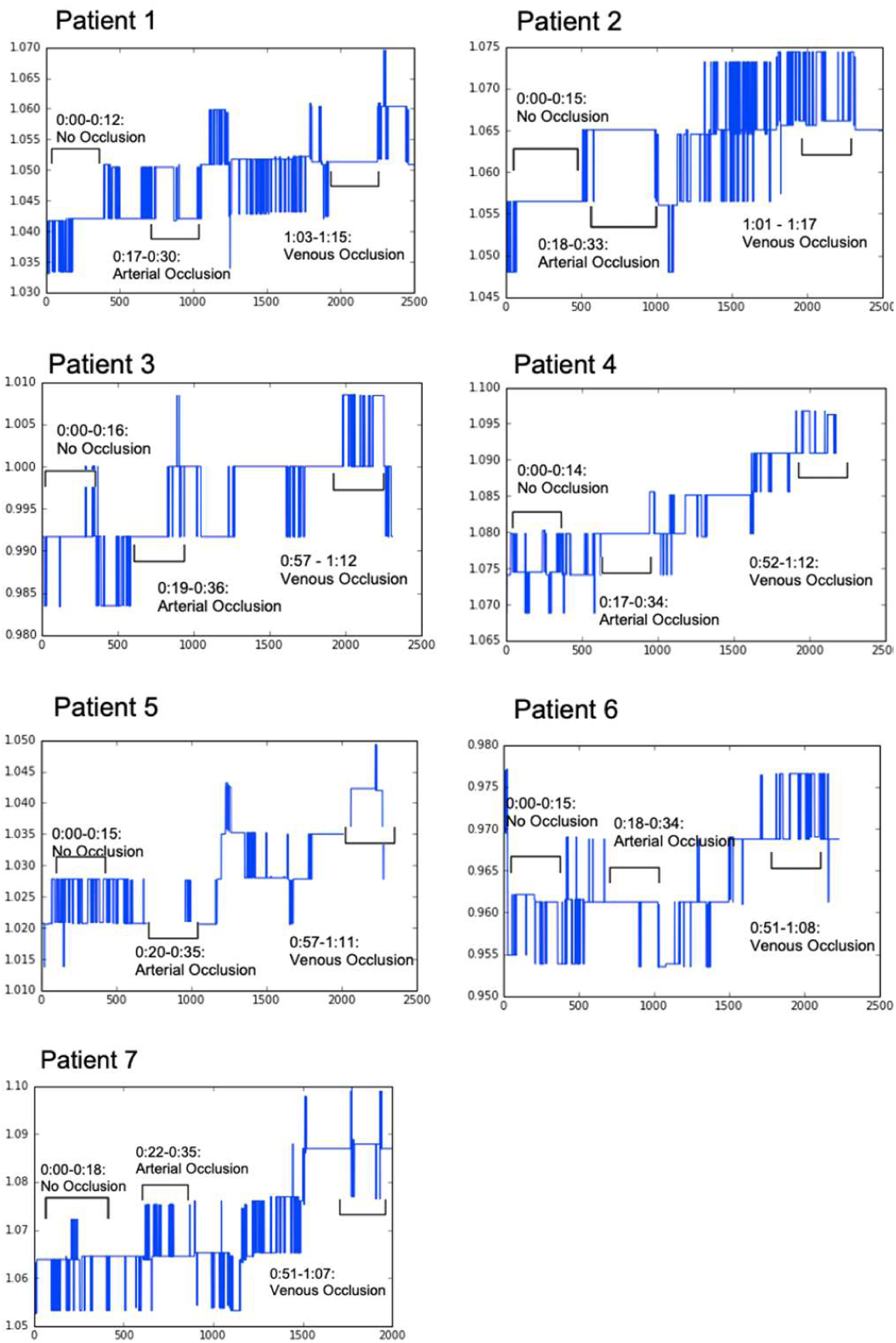
Output showcasing the “delta” between hands undergoing occlusion and a control hand. There was a distinct pattern for both an increase in arterial and venous occlusion across all 7 patients. X: Time - Represented by Frames in each Video (33 frames/ Second) Y: Magnitude of Change in Total Red Pixels (For Occluded Area/ Normal Control)

There was a consistent pattern amongst patients returning from arterial occlusion to no occlusion as well that consisted of an increase in rate of average pixel change oscillations and greater range (The lowest bound corresponding to each individuals no occlusion chart, and the highest bound corresponding to that for venous occlusion).

The change in average pixels per frame increased for arterial occlusion and even more so for venous occlusion.

**Figure 5:**
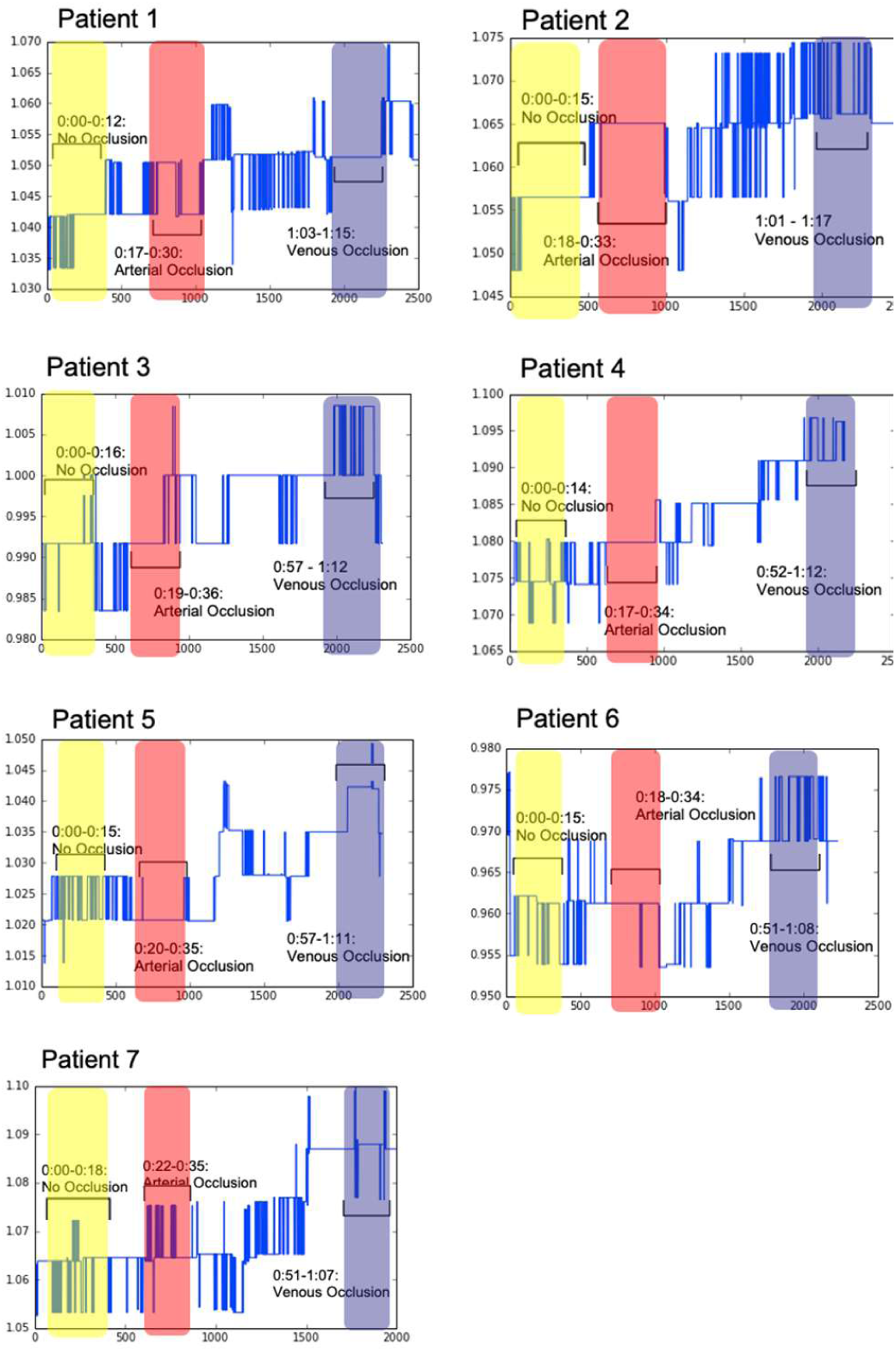
Output showcasing there was a distinct pattern for both an increase in arterial and venous occlusion across all 7 patients. There was a distinct pattern for both an increase in arterial and venous occlusion across all 7 patients. X: Time - Represented by Frames in each Video (33 frames/ Second) Y: Magnitude of Change in Total Red Pixels (For Occluded Area/ Normal Control)

## IV. Discussion

This pilot study showcased the feasibility of a free flap monitoring system using only a smart phone and machine learning. This solution was a considerably less computationally intense solution compared to other available methods and still was non-invasive. Any modern camera phone could theoretically download an app of this nature and produce accurate results to test both pulse or blood perfusion. The pulse detection system revealed that there is a noticeable however small delta in skin tone during a heart beat. Although this may be difficult to pick up by the human eye, it is potentially uncovered using pattern recognition and machine learning.

The free flap monitoring system was able to detect instances of no occlusion from arterial and venous occlusion by comparison against a control. The arterial occlusion showed less variability (5% increase) than venous occlusion (7% increase). Although noticeable changes also occurred in the other colors, due to the nature, consistent results, and ease of interpretation across patients the red pixel delta was chosen.

One potential reason for this change in pixels is the small change in skin tone that can occur from the act of bringing blood to the region. In arterial and venous occlusion, blood is either drained or congregated in the hand (respectively) which can lead to a more drastic and detectable change in skin tone. Using this technology would allow for a affordable, non-invasive, easy to use, and less computationally intensive method that could consistently flag instances of occlusion. One of the better benefits of using a pattern recognition algorithm is that it works by flagging a delta so this could be adapted for untrained labor to have a simple indicator flagging if a patient was potentially undergoing arterial or venous occlusion.

This study could be expanded by testing this technology in a non-simulated clinical setting. With more clinical data and earlier clinical data this technology could be potentially used to find automated measures that could predict occlusion before it occurs.

## V. Conclusion

Our smartphone video capture and analysis facilitates visualization of skin perfusion and can distinguish between states of no occlusion, arterial occlusion, and venous occlusion. The pattern shown after recovering from occlusion suggests that a similar pattern might be observed in tissue about to undergo occlusion, and implicates future studies could isolate diagnostic biomarkers before occlusion. This study shows promise for the use of inexpensive smartphone monitoring in a clinical setting for accurate free flap monitoring.

